# Analytical Validation of Automated DNA Isolation from Meat for High-Quality PCR-Based Food Authentication

**DOI:** 10.64898/2026.07.17.739293

**Authors:** Yulia K. Dewi, Yusrina Nabila C. Chudori

## Abstract

Reliable DNA isolation is a critical prerequisite for PCR-based food authentication, particularly for meat products where complex matrices may compromise DNA quality and amplification efficiency. This study aimed to analytically validate an automated DNA extraction method from meat matrices using Qiagen QIAcube Connect in combination with the DNeasy® Mericon Food Kit. Validation parameters included DNA concentration, total yield, purity, integrity, and assessment of PCR inhibitors using real-time PCR targeting the porcine *cytochrome b* gene. The method produced a mean DNA concentration of 219.5 ng/µL with an average yield of 21,519.7 ng, exceeding predefined acceptance criteria. Agarose gel electrophoresis confirmed DNA fragment sizes larger than the target amplicon, indicating suitability for PCR analysis. Real-time PCR evaluation demonstrated excellent linearity (R^2^ = 0.99–1.00), amplification efficiencies between 90.34% and 99.84%, and mean ΔCt values of 0.10, confirming the absence of PCR inhibition. These results indicate that the validated automated method is robust, reproducible, and suitable for routine PCR-based meat species authentication in food control laboratories.

---

Authentication of meat products has become increasingly important due to growing global concerns over food fraud, halal certification, and consumer safety [1][2]. Traditional morphological and protein-based methods often lack specificity, especially in processed foods where proteins degrade [5] [6]. In contrast, DNA-based methods, particularly real-time PCR, offer high specificity and sensitivity even in complex matrices [3] [1].

Real-time PCR has emerged as the gold-standard technique for species-level meat authentication due to its capacity for quantitative analysis, high analytical sensitivity, and resistance to complex matrix effects [8]. Real-time PCR enables the detection of trace DNA sequences from target species with excellent amplification efficiency and specificity, facilitating confirmation or exclusion of undeclared animal ingredients in meat products [8]. Despite the power of PCR-based assays, successful amplification critically depends on the quantity and quality of the input DNA. Meat and processed food matrices often present obstacles such as high fat content, protein-complex formation, and co-extraction of PCR inhibitors, which can impede downstream amplification [8] [1]. Efficient DNA extraction protocols that provide sufficient yield, purity, and fragment integrity are therefore essential to ensure reliable PCR results [1].

Automated DNA extraction platforms offer a promising solution to these challenges by minimizing operator variability, reducing contamination risk, and standardizing sample processing. While existing studies have focused on manual extraction methods, there is a critical need to validate automated workflows for routine application in food authentication laboratories. Recent literature suggests that incorporating automation with well-characterized extraction kits can improve reproducibility and allow accurate species identification across diverse meat matrices [2] [8] .

In this context, the present study aims to provide a comprehensive analytical validation of an automated DNA isolation method from meat matrices using the QIAcube Connect platform coupled with the DNeasy® Mericon Food Kit. By assessing DNA yield, purity, integrity, and PCR performance metrics, we aim to demonstrate the suitability of this approach for routine PCR-based meat species authentication.

## 1. Materials and Methods

### 2.1 Meat Matrix and Sample Preparation

Raw pork meat was used as the representative meat matrix in this validation study, considering its frequent use in food authentication research and its relevance in regulatory and halal verification contexts. Fresh meat samples were homogenized to ensure matrix uniformity prior to DNA extraction. Approximately 200 mg of homogenized sample was used for each extraction, consistent with internationally recommended practices for DNA-based food analysis [1] [8]. Sample homogenization is a critical pre-analytical step to minimize heterogeneity and improve DNA recovery from complex food matrices, particularly those with high protein and lipid content such as meat [2].

### 2.2 Automated DNA Extraction

DNA extraction was performed using the Qiagen QIAcube Connect automated platform in combination with the DNeasy® Mericon Food Kit, following the manufacturer’s instructions with minor adjustments tailored to meat matrices. The automated protocol “Small Fragments” was selected to optimize recovery of fragmented DNA typically encountered in food samples.

The extraction workflow included enzymatic lysis using proteinase K, organic phase separation to reduce protein and lipid contaminants, and silica membrane-based purification. DNA was eluted in a final volume of 100 µL elution buffer. Automation was employed to enhance reproducibility, reduce analyst-dependent variability, and minimize the risk of cross-contamination, which are critical factors in routine food testing laboratories [4] [7].

### 2.3 DNA Concentration and Purity Assessment

DNA concentration was determined by ultraviolet spectrophotometry at 260 nm. DNA purity was evaluated by calculating the absorbance ratio at A260/A280. These parameters were used to assess protein contamination and overall extract quality. Spectrophotometric analysis remains a widely accepted method for preliminary DNA quality assessment in food authentication studies, providing rapid and reproducible measurements suitable for routine laboratory workflows [4].

### 2.4 DNA Integrity Evaluation

DNA integrity was assessed using horizontal agarose gel electrophoresis (1% agarose, TAE buffer). Electrophoretic patterns were visually examined to evaluate fragmentation and to confirm that the average fragment size exceeded the length of the PCR target amplicon. Although food-derived DNA is frequently fragmented due to processing and matrix effects, DNA fragments larger than the target amplicon are considered sufficient for reliable PCR amplification [2] [8]. This evaluation step is therefore essential to confirm method suitability for downstream molecular analysis.

### 2.5 Assessment of PCR Inhibition

The presence of PCR inhibitors was evaluated using real-time PCR targeting the porcine *cytochrome b* gene. Serial dilutions of extracted DNA (0.5 to 0.00005 ng/µL) were prepared to generate standard curves. Amplification performance was assessed by calculating slope values, amplification efficiency, coefficient of determination (R^2^), and ΔCt values between extrapolated and measured Ct values. PCR inhibition was considered absent when amplification efficiencies ranged between 80–110%, R^2^ values exceeded 0.98, and ΔCt values were below 0.5, in accordance with internationally accepted validation criteria for PCR-based food analysis [3] [4].

### 2.6 Validation Parameters and Acceptance Criteria

Method validation focused on analytical performance parameters relevant to PCR-based food authentication, including DNA yield, repeatability, integrity, and PCR compatibility. Acceptance criteria were defined prior to experimentation and aligned with published guidelines and recent validation studies in food molecular analysis [3] [1].

## 3. Results and Discussion

### 3.1 DNA Yield and Concentration Performance

Automated DNA isolation method produced consistently high DNA concentrations across all validation runs. As summarized in Table 1, the mean DNA concentration reached 219.5 ng/µL, with an average total DNA yield of 21519.7 ng, exceeding the predefined acceptance criteria for PCR-based food authentication. Relative standard deviation (RSD) for DNA concentration (16.1%) and total yield (23.8%) reflects acceptable repeatability for biological matrices, where intrinsic heterogeneity is unavoidable. Comparable variability has been reported in recent validation studies involving raw and processed meat samples [1] [4]. The use of the automated extraction platform based on Qiagen QIAcube Connect likely contributed to improved consistency by reducing analyst-dependent variability and standardizing critical extraction steps.

**Table 1:** DNA concentration and yield obtained from automated extraction of meat matrices.

| Parameter | Mean | Standard Deviation | RSD (%) | Acceptance Criteria |
| --- | --- | --- | --- | --- |
| DNA concentration (ng/μL) | 219.5 | 35.4 | 16.1 | $\geq 4$ ng/μL |
| Total DNA yield (ng) | 21519.7 | 2749.5 | 23.8 | $\geq 200$ ng |

### 3.2 DNA Integrity Assessment

DNA integrity was evaluated by agarose gel electrophoresis. Representative electrophoretic profiles are shown in Figure 1. The extracted DNA exhibited fragmented patterns, which are typical for food-derived DNA, particularly from meat matrices. Importantly, the majority of DNA fragments were larger than the target PCR amplicon (>131 bp), indicating that the DNA was suitable for downstream real-time PCR amplification. Recent studies emphasize that fragmented DNA is not a limitation for PCR-based food authentication as long as fragment sizes exceed the target amplicon length [8]. Results obtained here are consistent with these findings and confirm the analytical fitness of the extracted DNA.

**Figure 1:**
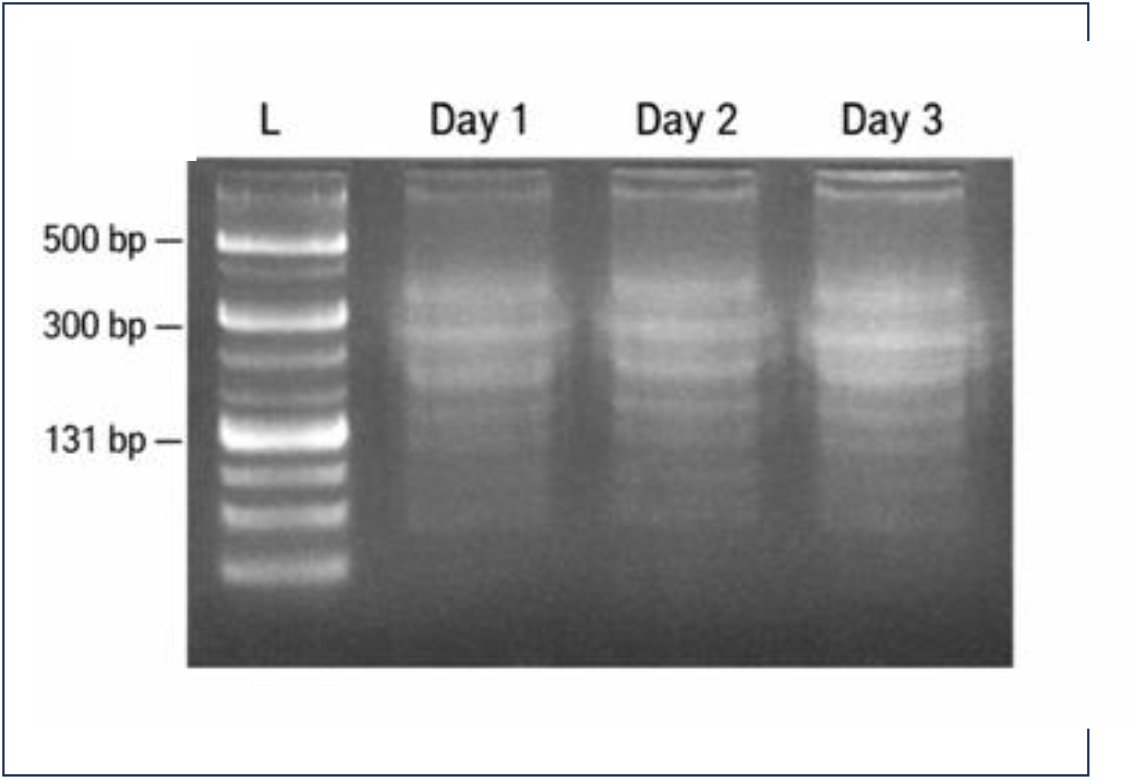
Agarose gel (1%) electrophoresis showing DNA extracted from meat samples over three validation days. Lanes show fragmented DNA with fragment sizes exceeding the target amplicon length (>131 bp), confirming suitability for PCR-based analysis.

### 3.3 PCR Inhibition and Amplification Efficiency

PCR performance was evaluated using real-time PCR targeting the porcine *cytochrome b* gene (*cytb)* across five serial DNA dilutions. Standard curves generated from these dilutions are summarized in Table 2, while representative amplification plots are shown in Figure 2. All standard curves demonstrated excellent linearity, with R^2^ values ranging from 0.99 to 1.00. Slope values (–3.3 to –3.6) corresponded to amplification efficiencies between 90.34% and 99.84%, which fall well within the optimal performance range defined by international validation guidelines [3]. Mean ΔCt value of 0.10 further confirmed the absence of significant PCR inhibition.

**Table 2:** Real-time PCR performance parameters for inhibitor evaluation (confirms robust PCR performance and absence of inhibitory effects in the extracted DNA). These performance parameters were evaluated in accordance with established quantitative PCR best-practice guidelines [10].

| Parameter | Range Obtained | Acceptance Criteria |
| --- | --- | --- |
| Slope | −3.3 to −3.6 | −3.1 to −3.6 |
| Amplification efficiency (%) | 90.34 – 99.84 | 80 – 110 |
| R<sup>2</sup> | 0.99 – 1.00 | > 0.98 |
| Mean ΔCt | 0.10 | < 0.5 |

**Figure 2:**
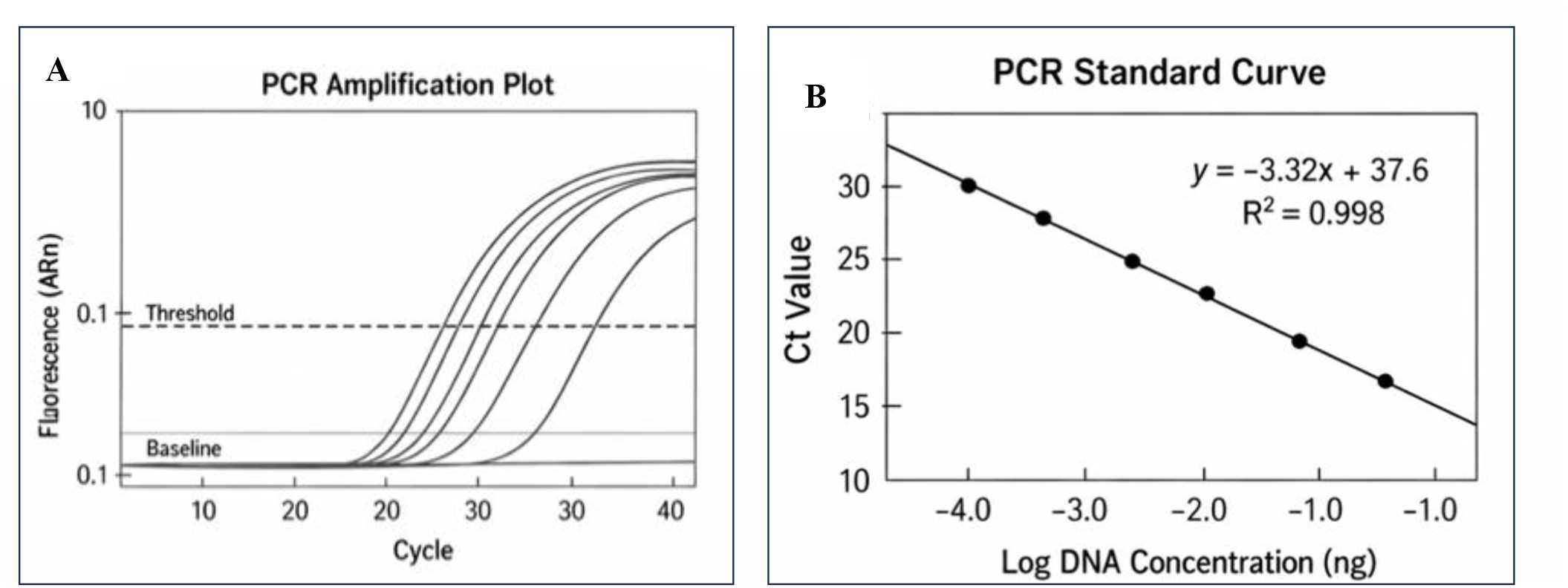
Representative real-time PCR amplification plots (A) and corresponding standard curves (B) obtained from serial dilutions of extracted DNA targeting the porcine *cytochrome b* gene. The linearity and amplification efficiency indicate absence of PCR inhibitors.

### 3.4 Implications for Routine Food Authentication

Combined results of DNA yield, integrity, and PCR performance demonstrate that the validated automated extraction method is robust and fit for routine PCR-based meat authentication. Integration of automation minimizes operator variability and supports standardized analytical workflows, which are increasingly required in regulatory and food control laboratories [1].

## Conclusions

This study provides a comprehensive analytical validation of an automated DNA isolation workflow for meat matrices intended for PCR-based food authentication. Validated method consistently produced high DNA yields with acceptable repeatability, preserved DNA fragment integrity compatible with real-time PCR requirements, and demonstrated robust amplification performance without detectable PCR inhibition. Across all validation parameters, including DNA concentration, total yield, fragment size distribution, amplification efficiency, linearity, and ΔCt values, the automated workflow fulfilled or exceeded internationally accepted acceptance criteria for molecular food analysis. The results confirm that fragmented DNA obtained from meat matrices remains analytically fit for PCR-based species identification when fragment sizes exceed the target amplicon length. The integration of automation into the DNA extraction process offers clear advantages over manual approaches, particularly in terms of reproducibility, standardization, and reduced risk of operator-dependent variability. These attributes are critical for routine application in food control, regulatory enforcement, and halal verification laboratories, where analytical reliability and method robustness are essential. Future studies should extend this validated workflow to mixed-species and highly processed meat products, as well as explore its integration with quantitative PCR and digital PCR platforms for enhanced regulatory enforcement.

In conclusion, the automated DNA isolation method validated in this study is suitable for high-quality PCR-based meat authentication and provides a reliable foundation for routine implementation in analytical laboratories. Future studies may extend this validation framework to additional meat species, processed food matrices, and quantitative adulteration scenarios to further support broader regulatory and industrial applications.

## Acknowledgements

This work was supported by institutional laboratory facilities. Authors would like to acknowledge the technical support and laboratory facilities that made this study possible. Appreciation is extended to all personnel involved in sample preparation, analytical measurements, and data acquisition. Authors also thank colleagues who provided constructive feedback during the development of this manuscript.

## References

[1] Adenuga, B. M., & Montowska, M. (2023). DNA-based methods for meat species authentication: Advances, challenges, and future perspectives. Comprehensive Reviews in Food Science and Food Safety, 22(3), 1764–1785. 10.1111/1541-4337.13142

[2] Azad, M. A. K., Dey, M., Khanam, F., Biswas, B., & Akhter, S. (2023). Authentication of meat and meat products using molecular assays: A review. Journal of Agriculture and Food Research, 12, 100586. 10.1016/j.jafr.2023.100586

[3] Broeders, S., Huber, I., Grohmann, L., Berben, G., Taverniers, I., Mazzara, M., Roosens, N., & Morisset, D. (2021). Guidelines for validation of qualitative real-time PCR methods for food and feed analysis. Trends in Food Science & Technology, 115, 229–241. 10.1016/j.tifs.2021.06.015

[4] Costa, J., Amaral, J. S., & Oliveira, M. B. P. P. (2022). Advances in DNA-based methods for food authenticity testing. Food Control, 132, 108527. 10.1016/j.foodcont.2021.108527

[5] Hu, J., Wei, H., Jiang, Y., Xue, Q., & Wang, F. (2025). DNA barcoding in meat authentication: Principles, applications, and future perspectives. Foods, 14(20), 3522. 10.3390/foods14203522

[6] Kim, Y., Lee, H.-S., & Lee, K.-G. (2022). Detection of porcine DNA in processed foods using real-time PCR. Food Science and Biotechnology, 32(1), 21–26. 10.1007/s10068-022-01169-x

[7] Mieczkowska, A., Kaczmarek, A., & Rybicka, I. (2021). Automation in DNA extraction workflows: Impacts on reproducibility and analytical performance. Analytical and Bioanalytical Chemistry, 413(18), 4503–4514. 10.1007/s00216-021-03421-5

[8] Minoudi, S., Pardo, M. A., García, T., & González, I. (2024). DNA-based approaches for meat species identification and quantification: Recent developments and analytical challenges. Food Chemistry, 431, 137098. 10.1016/j.foodchem.2023.137098

[9] Muflihah, A., Hardianto, A., Kusumaningtyas, P., Prabowo, S., & Hartati, Y. W. (2023). DNA-based detection of pork adulteration in food products: Analytical performance and application. Heliyon, 9(3), e14418. 10.1016/j.heliyon.2023.e14418

[10] S. A. Bustin, V. Benes, T. Nolan, and M. W. Pfaffl, “Quantitative real-time RT-PCR – a perspective,” Journal of Molecular Endocrinology, vol. 34, no. 3, pp. 597–601, 2009, doi: 10.1677/JME-04-0025

[11] Saleem, A., Imtiaz, A., Yaqoob, S., Awais, M., Awan, K. A., Naveed, H., Khalifa, I., Ercisli, S., & Mugabi, R. (2025). Integration of conventional and molecular systems for meat authentication: A review with bibliometric analysis. Food Chemistry X, 29, 102778. 10.1016/j.fochx.2025.102778

